# Chromatin maturation of the HIV-1 provirus in primary resting CD4^+^ T cells

**DOI:** 10.1101/526053

**Authors:** B Lindqvist, S Svensson Akusjärvi, A Sönnerborg, M Dimitriou, JP Svensson

## Abstract

Human immunodeficiency virus type 1 (HIV-1) infection is a chronic condition, where viral DNA integrates into the genome. Latently infected cells form a persistent, heterogeneous reservoir. The reservoir that reinstates an active replication comprises only cells with intact provirus that can be reactivated.

We confirmed that latently infected cells from patients exhibited active transcription throughout the provirus. To find transcriptional determinants, we characterized the establishment and maintenance of viral latency during proviral chromatin maturation in cultures of primary CD4^+^ T-cells for four months after *ex vivo* HIV-1 infection. As heterochromatin (marked with H3K9me3 or H3K27me3) gradually stabilized, the provirus became less accessible with reduced activation potential. In a subset of infected cells, active marks (i.e., H3K27ac) remained detectable, even after prolonged proviral silencing. After T-cell activation, the proviral activation occurred uniquely in cells with H3K27ac-marked proviruses. Our observations suggested that, after transient proviral activation, cells were actively returned to latency.

**Author Summary:** HIV infection is a devastating disease affecting 35 million people worldwide. Current anti-retroviral treatment is highly effective and has made the HIV infection chronic. However, a cure is far on the horizon. The problem for an HIV cure is that, even though the virus particles are eradicated, the infected cells maintain the information of remake the virus. This information is integrated in the host cell. The proviral chromatin switches between active and inactive states. Thereby, the infected cells evade both the immune system and death associated with massive viral production.

Here we have characterized the composition of the proviral chromatin and how it connects with transcription and viral production. In resting primary CD4^+^ T-cells, we follow the fate of the provirus starting at infection until latency is firmly established. We found that only a fraction of the proviruses switched between the two chromatin states, and thus were able to remain undetectable but still produce new viruses. These proviruses were associated with a specific chromatin mark, allowing us to identify the activatable fraction of infected cells. Our study provides key insights as to detect the remaining HIV-infected cells capable of reseeding the infection and the mechanisms whereby these cells are maintained.

## Introduction

Once human immunodeficiency virus type 1 (HIV-1) infects a cell, typically an activated CD4^+^ T-lymphocyte, the most likely outcome is cell death by apoptosis or pyroptosis [1,2]. In some cells, the viral genome integrates into the host chromosome. Upon integration, the viral sequence is packaged into chromatin, and in a subset of cells, as the cell returns to quiescence, the proviral chromatin then becomes condensed and silenced together with large portions of host chromatin [3]. At any time can the provirus reactivate, leading to viral gene expression and virus production that may cause cell death. The resulting reservoir of rare latently infected cells (∼1 in 10^6^ CD4^+^ T-cells) is the main obstacle to finding a cure for HIV-1/AIDS.

Proviral latency occurs very early [4,5] as the reservoir is established within days of the initial infection [6-8]. After residual viral proteins are degraded, latently infected cells are indistinguishable from uninfected cells [9]. Consequently, the virus escapes the immune system and the actions of current drugs. However, functional, intact proviruses can recommence the replication when latency is reversed. A means for identifying this functional reservoir would represent a main milestone in clinical advances.

The latent reservoir displays heterogeneity in the types of cells infected, the anatomical location in the body, and the strength of silencing. Also, the reservoir harbors only a small fraction (5-12%) of functional, intact proviruses [10-14]. The main fraction of proviruses are defective; they contain large internal deletions, which occur through template-switching during reverse transcription [15], or they contain G-to-A mutations, induced by apolipoprotein B mRNA editing enzyme (APOBEC) catalytic subunit 3G [16]. In time, the viral reservoir in a patient evolves. Even though the defective proviruses cannot produce infective particles, they can act as decoys to the immune system, and thus, they shape the reservoir [17]. Moreover, the detection of HIV-1 particles by CD8^+^ cells does not necessarily lead to HIV-1 elimination [18]. In fact, it may lead to proliferation and clonal expansion of the reservoir, which suggests that either the provirus can maintain its latent state, when the host cell is activated, or it can be actively resilenced. Reactivated latently infected T-cells have developed mechanisms that may allow cell division without activating virally-induced cell death pathways [19].

The activation potential of the provirus depends on several factors, including the epigenetic context and the nuclear environment [20-23]. HIV-1 is guided to active regions by the viral integrase, which has a high affinity for the host cellular replication cofactor, lens epithelium-derived growth factor (LEDGF). LEDGF recognizes histone H3 trimethylated at lysine 3 (H3K36me3) and H3K4me1 [21]. During T-cell activation, the provirus becomes anchored to the nuclear pore near open chromatin domains [24]. Among productively infected cells and reactivated latently-infected cells in the reservoir, proviruses are mainly found in generally active or poised chromatin; in contrast, permanently silenced proviruses are found in regions of heterochromatin [20]. Proviral integration into cell cycle genes appear to be more reactivatable and are subject to spontaneous reactivation [25]. Contrary to the expectation that cells with reactivated proviruses would be cleared by the immune system, patients that received long-term anti-retroviral therapy (ART) showed enrichments of clones with HIV-1 integrations into genes associated with proliferation and survival [26,27].

Although the HIV-1 provirus is integrated into active chromatin, as infected cells return to quiescence, the inactive proviral chromatin often becomes associated with repressive histone modifications, such as the facultative heterochromatin mark, H3K27me3, or the constitutive heterochromatin mark, H3K9me3 [28-32]. Accordingly, polycomb repressive complex 2 (PRC2) and euchromatic histone-lysine N-methyltransferase 2 (EHMT2) are required for the establishment and maintenance of HIV-1 proviral silencing [33]. A lack of H3K27me3 and H3K9me2/3 sensitizes latent proviruses to latency reversal agents (LRAs) [33-35]. The process of establishing heterochromatin is lengthy and complex [36]. Upon HIV-1 infection, different epigenetic marks are initially established over the provirus, but the H3K27me3-to-H3K9me3 ratio (H3K27m3/H3K9m3) evolves over time [37]. Due to the compact nature of heterochromatin structures, access to the transcriptional machinery at the canonical long-terminal repeat (LTR) promoter is restricted, and thus, HIV-1 transcription is hampered and HIV-1 latency is promoted (Tyagi and Karn, 2007; Tyagi et al., 2010). However, RNA polymerase II (RNAPII) and active chromatin marks, such as H3K4me3, have been shown to remain on the LTR promoter, apparently to maintain the promoter in a state poised for transcription. Short transcripts from the promoter-proximal trans-activation response element (TAR) have been identified in latently infected cells [38]. The bromodomain and extra-terminal domain (BET) protein, BRD4, is present at the latent provirus. Removal of BRD4 by the BET inhibitor, JQ1, leads to the release of RNAPII proximal-pausing [39]. The post-initiation block of RNAPII has long been recognized as a rate-limiting step of latency reversal. However, recent data have also highlighted the roles of blocks in elongation, splicing, and termination. These major HIV-1 transcription-restrictive factors are important, as transcripts from latently HIV-1 infected cells in patients arise from points along the entire provirus [40].

A strong stimulus for reversing proviral latency is the activation of T-cell receptors (TCRs). However, only a few infected cells (<5%) display proviral activation upon a single round of TCR activation [20]. Both TCR activation and LRA administration have been shown to elevate viral RNA levels, but these treatments have modest effects on reduction of the latent reservoir HIV-1 patients [41-45].

Here, we dissected the chromatin and RNA landscape of the HIV-1 provirus during latency establishment in primary resting CD4^+^ T-cells. We aimed to reveal the mechanisms that maintained the activation potential of latently infected cells.

## Results

### Transcription over the entire provirus isolated from patient cells

To confirm that the entire provirus was transcribed in cells from patients successfully treated with ART, as suggested previously [40], we isolated RNA from peripheral blood mononuclear cells (PBMCs) from five patients with HIV-1 infection of diverse subtypes. These five patients responded well to treatment with virus levels <50 copies/µl for a median of 8 years (range 1.8–20 years) and increase of CD4^+^ T-cells to levels within the normal limits (Table S1). The levels of cell-associated (CA) RNA were measured with reverse transcription (RT), followed by droplet digital PCR (ddPCR). Several primer pairs were designed over the proviral genome (Fig S1). In unstimulated PBMCs, viral CA-RNA levels were significantly (p>0.05) higher than background (*read-through*) levels of transcription (Fig 1A). In most patients, the highest RNA levels originated from the TAR region of HIV-1. The abundance of mature multiply-spliced transcripts (*tat-rev*) was significantly lower (p<0.05) than the abundances of five unspliced products. This indicated that latent proviruses were actively transcribed to a low degree, but still, mature transcripts failed to emerge.

**Fig 1.**
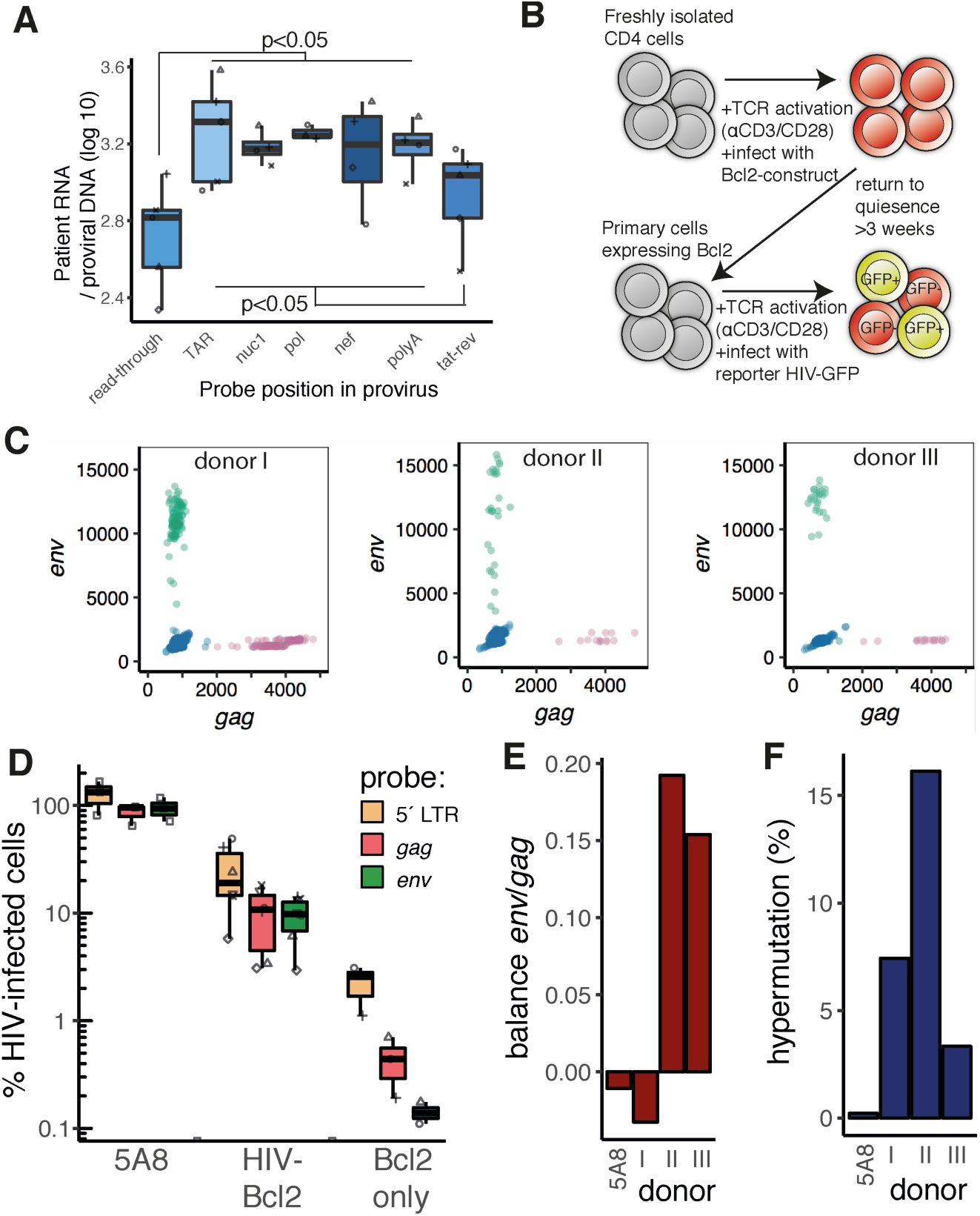
Transcripts in HIV-1 patient cells and infection of primary T cells. (**A**) Box plot showing transcription levels in PBMCs from well-treated patients (*n*=5), at various positions of the provirus. Each patient is represented by a unique symbol. (**B**) Schematic of the generation of HIV-1 carrying Bcl2 model cells. (**C**). ddPCR plots of HIV-1 infected Bcl2-cells from three healthy donors with probes against *env* (y-axis) and *gag* (x-axis). DNA was isolated at 3 dpi. (**D**) Quantification of ddPCR results over three probes (*5’LTR, gag* and *env*). (**E**) Ratios of *gag*/*env* ddPCR signals to reflect imbalances between 5’ and internal large deletions in the three donors. (**F**) Fraction of “rain”, i.e. low *env* signals (<10,000 a.u.) reflecting APOBEC3G-induced hypermutations.

### An established primary cell model with a single-round reporter

To elucidate the underlying transcriptional and chromatin landscape, we turned to an established model for HIV-1 latency in primary human CD4^+^ T cells. Technical and ethical issues hindered molecular characterization of chromatin in patient cells. This Bcl2-model comprised CD4^+^ lymphocytes isolated from healthy individuals, which are amenable to culture for extended periods of time [46,47] (Fig 1B). Here, we isolated CD4^+^ cells from fresh peripheral blood collected from three healthy donors. These cells were immediately stimulated with antibodies against CD3 and CD28 (αCD3-CD28) and grown in media containing growth factors. After 72 h, cells were transduced with a lentiviral vector that carried the gene that encoded anti-apoptotic Bcl2. Cells were then returned to quiescence by culturing in cytokine-free media. After three weeks, the cells were either cryopreserved or re-stimulated with αCD3-CD28 for 72 h. Activated cells were infected with a HIV-1 reporter virus and maintained under stimulating growth conditions for an additional three days. The HIV-1 reporter virus was rendered replication-deficient with six mutations, but it encoded the full-length viral genome. In addition, the *GFP* gene was inserted in the *env* coding region. Bcl2-model cells isolated from the three healthy individuals were divided into two groups; one was infected with HIV-1 immediately, and the other was infected after one freeze-thaw cycle.

To determine the fraction of initially infected cells, we isolated DNA at three days post infection (dpi). We quantified the levels of proviral DNA with ddPCR and three probes that targeted *gag* and *env* (Fig 1C), as well as the 5’ LTR [48,49]. For comparison, we included the J-lat clone 5A8, a Jurkat-derived cell line with one integrated latent HIV-1 reporter provirus per cell [22,50]. Among the Bcl2-model cells, the percentage of successful HIV-1 infections ranged from 3.1% to 18.2% (median: 10.8%), estimated with the standard *gag* probe, in parallel with the *env* probe (median: 9.8%) (Fig 1D).

The primary cells were infected with two lentiviruses, the Bcl2 construct and the HIV-1 construct, both containing a 5’ LTR sequence; thus, the 5’ LTR probe signal was detectable at high levels (median: 19.5%). After the Bcl2 transduction, most model cells remained in a primary state and were not transformed, as the 5’ LTR signal showed that only a minority of the model cells contained the Bcl2 construct. This finding suggested that, rather than protecting individual cells from apoptosis, the Bcl2-infected cells likely acted as isogenic feeder-cells that secreted factors to sustain healthy primary cultures. The lentiviruses were nearly exclusively integrated, because the 2-LTR circles were observed at low or undetectable levels (Fig S1).

### Intact provirus in a subset of cells

To determine the initial fraction of cells with intact proviruses, we quantified the prevalence of proviruses with large internal deletions and APOBEC3G-induced hypermutations. These mutational events typically occur before or during the integration of viral DNA.

Large deletions were estimated by the imbalance between the 5’ *gag* and internal *env* signals (Fig 1E); this imbalance was expressed as the log2 of the *env/gag* ratio. The donors showed large variations, ranging from log2 (*env/gag*) of −0.03 to +0.19. As expected. DNA from the control 5A8 cells, with a single intact provirus per cell, displayed a uniform ddPCR signal across the provirus (log2 (*env*/*gag*) = −0.01).

To calculate the fraction of hypermutated proviruses, the *env* probe had been designed to target an APOBEC3G hotspot. The primers and probe matched a cluster of 12 previously described APOBEC3G-induced mutations [51]. The APOBEC3G-induced G-to-A mutations reduced the efficiency of the PCR reaction, and thus, they produced a lower signal in a droplet containing a single mutated template. This result was reflected in the characteristic “rain” signal observed below the cluster of *env*^+^ droplets (Fig 1C, most pronounced in donor II; Table S2). The donor cells showed varying levels of hypermutations, ranging from 3.3% to 16% (Fig 1F). The 5A8 cells were used to confirm that virtually no “rain” (0.2%) signal was detected in settings without hypermutations. These results confirm that the large majority of the proviruses in the model cells are intact.

### Number of HIV-1 infected cells diminishes in time

At 72 h after HIV-1 infection, we returned the cells to the resting state by transferring them to cytokine-free media (Fig 2A). The cells were maintained in culture and followed for four months. Samples were collected at 30, 70, 90, and 120 dpi. To determine the number of HIV-1 infected cells in the total culture, we quantified the fraction of cells that harbored the provirus at each time point. We followed the decline of HIV-1 infected cells, as they were competed out by uninfected cells, by performing ddPCR on genomic DNA. We used *gag* and *env* probes to identify HIV-1 unique regions (Fig 2B), and 5’ LTR probes to detect both the HIV-1 and the integrated Bcl2 segment (Fig 2C). At 90 dpi, the estimated fraction of HIV-1 infected cells was 2.2% (median *gag*) or 1.6% (median *env*). The 5’ LTR signal leveled out over time; at 90 dpi, the signal indicated 9.0% (median) remaining infected cells, which suggested approximately 7% of cells containing the *bcl2* gene. The resilience of the 5’ LTR signal might reflect the survival advantage conferred by the Bcl2 protein in long-term cell cultures. These data show that the cultures are not stable, but continue to evolve after the initial latency establishment.

**Fig 2.**
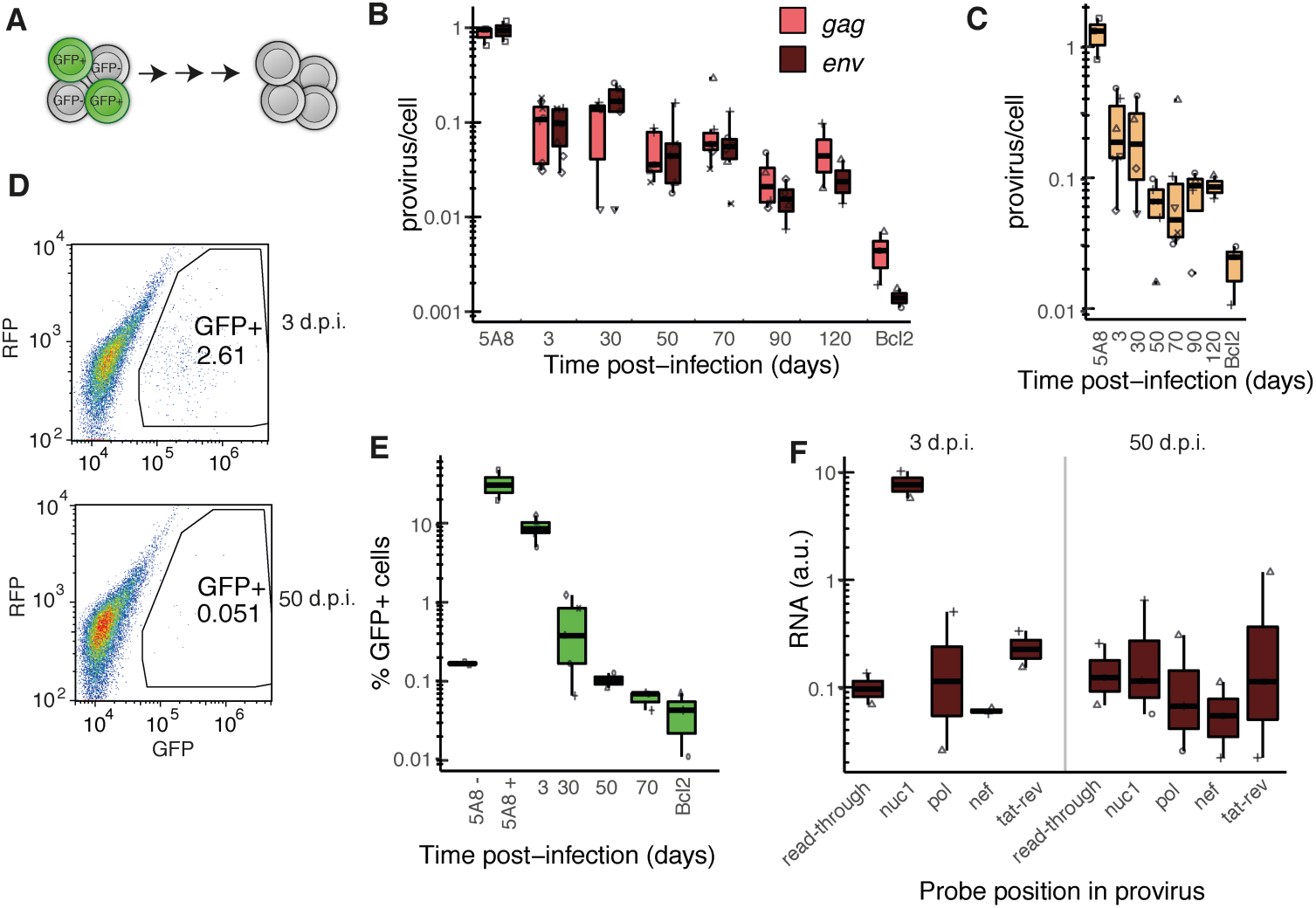
Fate of the HIV-1 provirus in resting T-cells in time. (**A**) Active cells were transferred to cytokine-free media to return to quiescence. (**B**) Proviruses quantified by ddPCR at the *gag* and *env* loci. Signals normalized to endogenous *rpp30*. J-lat clone 5A8 has a single integrated provirus and the Bcl2-only cells have been infected with the Bcl2-virus but not the HIV-reporter virus. (**C**) Proviruses quantified by ddPCR at the *5’ LTR* locus. Signals normalized to endogenous *rrp30*. (**D**) Flow analysis of cells at 3 and 50 dpi gated for GFP-positive cells expressing the GFP-containing HIV-reporter provirus. (**E**) Quantfication of (D) in time. (**F**) RT-ddPCR results of cell-associated RNA isolated at 3 and 50 dpi. Probes were designed along the provirus (*read-through* before transcription start at the LTR promoter), *tat-rev* recognizes the multiple-spliced transcript.

### Spontaneous HIV-1 activity

Next, we determined the prevalence of productively infected cells over time. First, GFP-positive cells in unperturbed cultures were counted with flow cytometry (Fig 2D-E). Here, 2– 6% of cells were initially GFP^+^ at 3 dpi (Fig 2E). Then, the GFP frequency rapidly declined and became indistinguishable from the GFP-negative control Bcl2-model cells at 70 dpi.

Second, as an alternative method for tracking provirus activity, we measured RNA levels in unperturbed cells. CA-RNA was isolated from cells at 3 and 50 dpi (Fig 2F). The RT-ddPCR results provided a population average, in contrast to flow cytometry data, which gave information on individual cells. Unlike cells from patients with HIV-1 (Fig 1A), in Bcl2-model cells infected with HIV-1, we could not capture TAR HIV-1 transcripts, which represented initial RNAPII products, because the sequence was identical between HIV-1 and Bcl2 constructs. Transcription through the first nucleosome (*nuc1*), which lies downstream of the TAR region, corresponds to early transcription extension. We found a significant *nuc1* RNA signal at 3 dpi. At 50 dpi, possibly for technical reasons, few data points were above the detection limit, as estimated by the signal from the read-through probe.

### Prolonged quiescence diminished proviral reactivation potential

To examine the mechanisms underlying proviral activation, we activated T cells with TCR stimulation. We stimulated the cells for 48 h with either αCD3-CD28, or phorbol 12-myristate 13-acetate (PMA), in conjunction with ionomycin (Fig 3A). T-cell activation was performed in the presence of raltegravir to hinder viral reintegration. The T-cell activation was phenotypically assessed and confirmed by cell growth, cell lumpiness, rapid media turnover, and increased cell death. Unexpectedly, these clearly activated cells largely failed to present the expected surface markers (CD25 or CD69) indicative of T-cell activation (Fig S3). The control J-lat 5A8 cells presented CD25 or CD69 after activation, which ruled out technical difficulties.

**Fig 3.**
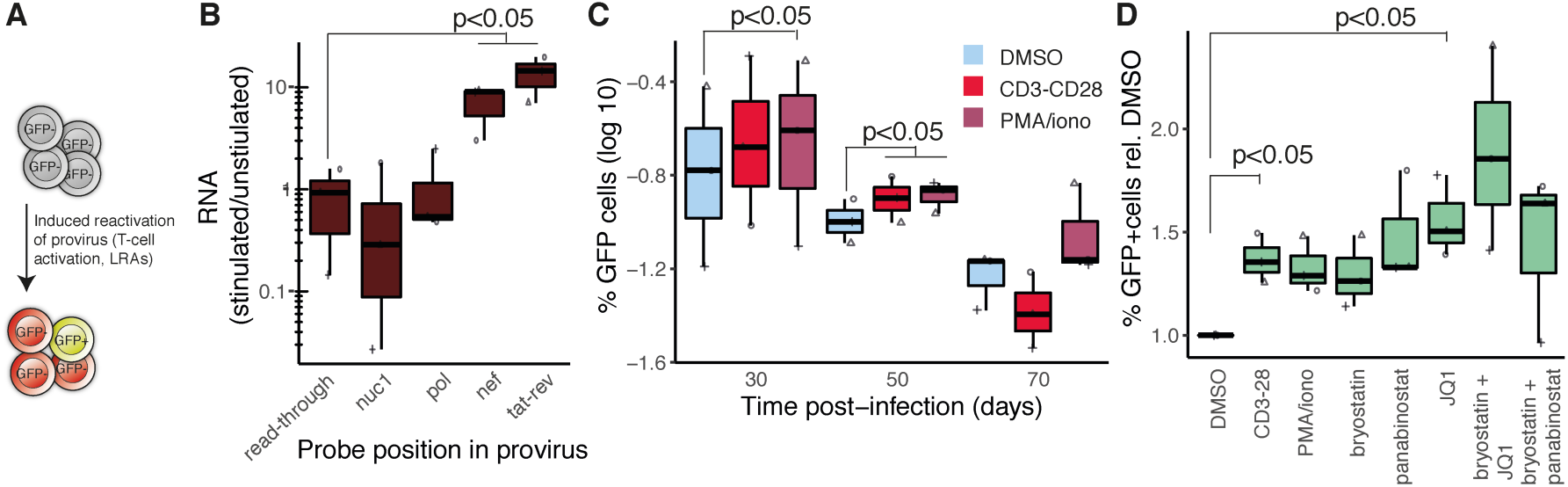
HIV-1 latency reversal by T-cell activation and LRAs. (**A**) Resting cells were treated with αCD3-CD28, PMA/ionomycin, or LRAs. (**B**) RT-ddPCR results at 4 proviral loci and the splice product *tat-rev* in activated/resting cells. (**C**) GFP-positive cells gated in flow cytometry at 30, 50 and 70 dpi after 48h treatment with DMSO, αCD3-CD28, or PMA/ionomycin. (**D**) GFP-positive cells gated in flow cytometry at 50 dpi after 48h treatment with LRAs.

To detect latent HIV-1 reversal, we again performed ultrasensitive CA-RNA quantification with ddPCR. At 50 dpi, the mature multiply-spliced transcripts were 10±3 fold increased (p<0.05) after 48 h of TCR stimulation (Fig 3B). The *nef* probe, which detected all transcripts completed to the 3’ end, showed a similar increase in transcription (9±3, p<0.05). The probes at the 5’ region did not detect an increase after TCR activation, consistent with an ongoing, non-productive transcription at the 5’ region [40]. Also, the read-through transcripts were not affected by T-cell activation, contesting an unspecific shift in chromatin accessibility.

To determine the number of cells with activated provirus, GFP-positive cells were detected with flow cytometry at three time-points (3, 30, and 50 dpi; Fig 3C). T-cell activation by both αCD3-CD28 conjugation and PMA/ionomycin resulted in a small, but significant increase (p<0.05) in cells with activated provirus, compared to untreated cells, but only under certain conditions.

Next, at 50 dpi, we exposed cells for 48h to a panel of previously described LRAs, including the HDAC inhibitor panabinostat, the protein kinase C agonist bryostatin, and the BET inhibitor JQ1. These drugs were administered in single regimens or in combinations, and in the presence of raltegravir to hinder viral reintegration. Cell viability was determined and only PMA/ionomycin had a significant (p<0.05) effect (Fig S4). The latency reversal results were unconvincing; only JQ1 alone or together with bryostatin could consistently induce proviral activation (Fig 3D). This implies that BET proteins, such histone acetyltransferase BRD4 [52], played a role in latency reversal in our primary CD4^+^ cells.

### RNAPII recruitment to proviral chromatin

We then aimed to relate the reactivation potential to the transcription machinery and the chromatin microenvironment. Previous studies have identified RNAPII at the inactive LTR promoter [38]. Here, we measured RNAPII at the provirus during the establishment of latency with chromatin immunoprecipitation, followed by quantitative PCR (ChIP-qPCR). To prevent erroneous signal from dead cells, prior to chromatin isolation viable cells were isolated by Ficoll density centrifugation. Two different forms of the RNAPII complex were investigated: the initiated RNAPII, detected by phosphorylated serine 5 (ser5ph) in the repetitive C-terminal domain of the RPT1 subunit (Fig 4A); and the elongating RNAPII, detected by phosphorylated serine 2 (ser2ph) in the same domain (Fig 4B). As before, we used J-lat clone 5A8 as a reference. Both forms of RNAPII were found at the site of the latent provirus at 30 dpi, and they remained at that site throughout the experiment, but the levels gradually diminished.

**Fig 4.**
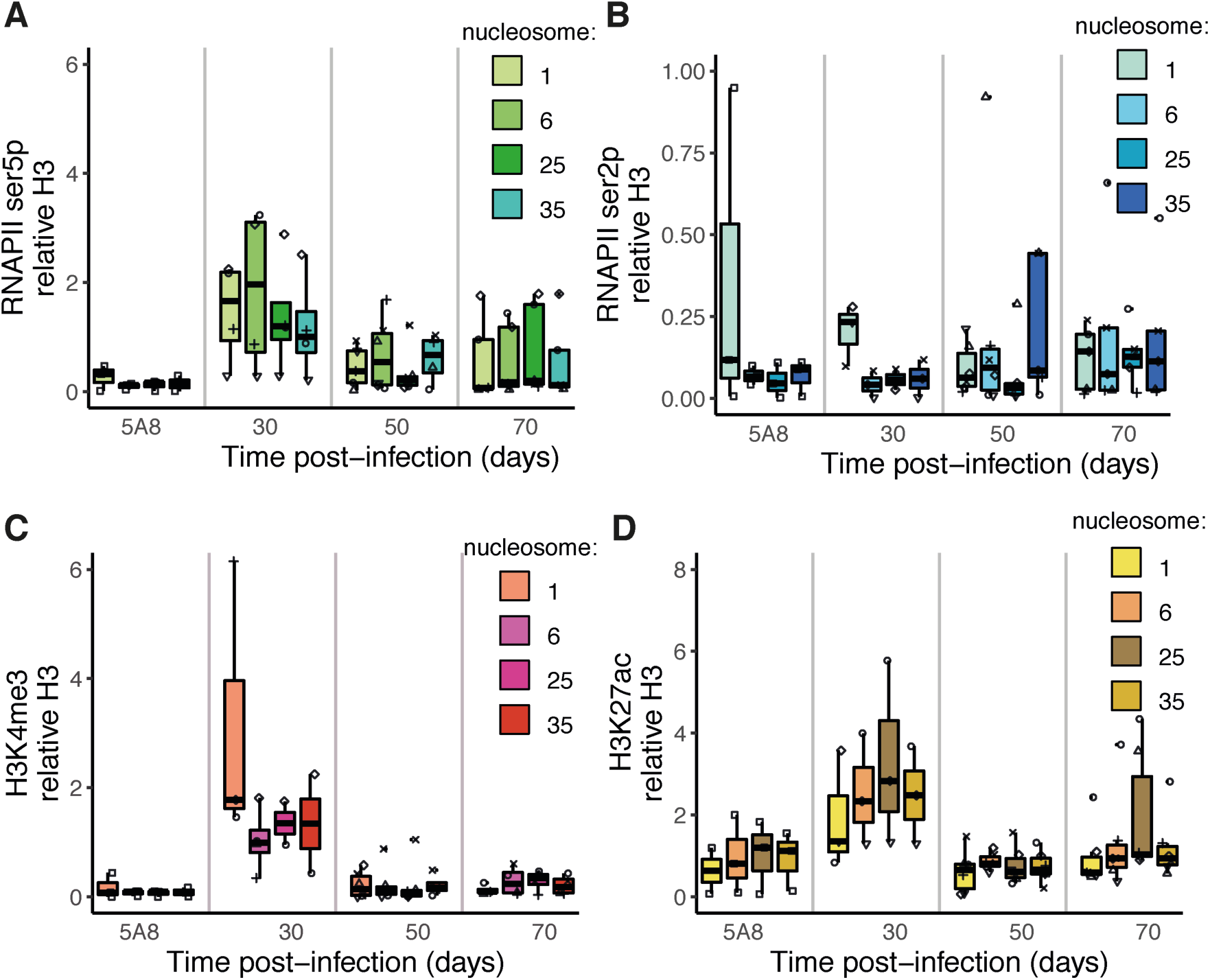
Active marks along HIV-1 proviral chromatin. (**A**) ChIP-qPCR signal for initiated RNA Pol II (CTD ser5 phosphorylated). (**B**) ChIP-qPCR signal for elongating RNA Pol II (CTD ser2 phosphorylated). (**C**) ChIP-qPCR signal for promoter mark H3K4me3 **D.** ChIP-qPCR signal for enhancer mark H3K27ac. Boxplots show data of T-cells from health donors (*n*=3, in duplicate).

### Active chromatin marks remain on the provirus

Upon integration, the proviral DNA sequence is rapidly encapsulated in chromatin. To determine the chromatin profile of the provirus, we followed the appearance of an array of chromatin marks. In our primary cells, the promoter mark, H3K4me3, was associated with the LTR promoter at early latency, but then, its abundance declined (Fig 4C). Another mark, H3K27ac, which associates with active enhancers and promoters, was found throughout the provirus life cycle, and its signal weakened at a rate similar to that of the other active marks, except that a low H3K27ac-signal was consistently detected (Fig 4D).

### Inactive chromatin marks accumulate

To follow the epigenetic inactivation of the provirus, we assessed the distribution of constitutive and facultative heterochromatin. The constitutive H3K9me3-mark appeared to be uniformly distributed over the proviral body by 50 dpi (Fig 5A). In contrast, the facultative H3K27me3-mark gradually became more prominent throughout the time-course (Fig 5B). In unstimulated J-lat 5A8 cells, both H3K9me3 and H3K27me3 were detected at relatively low levels in the proviral body (i.e., excluding the LTR promoter).

**Fig 5.**
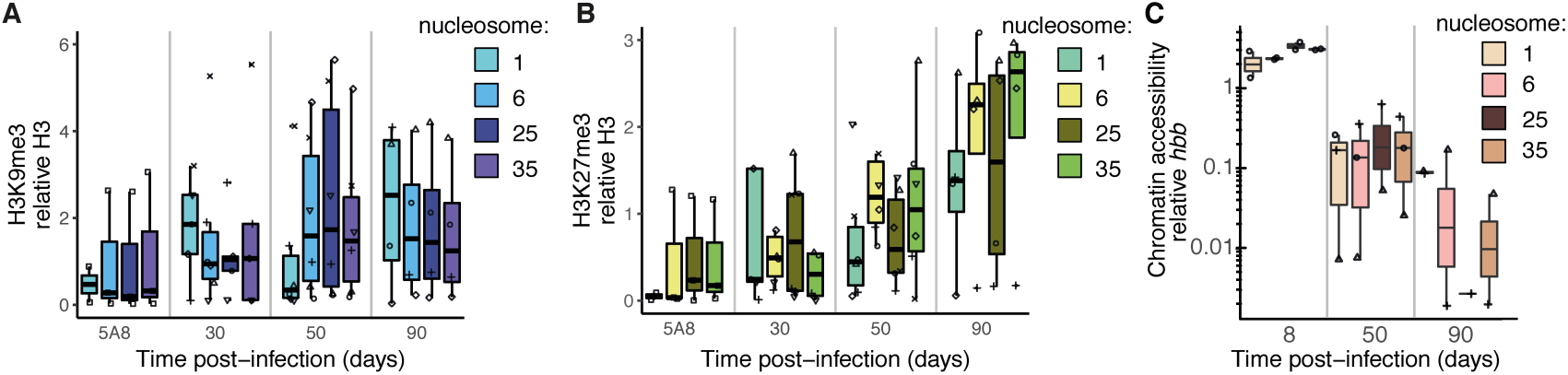
Heterochromatin and chromatin compaction along HIV-1 proviral chromatin. (**A**) ChIP-qPCR signal for constitutive heterochromatin mark H3K9me3. (**B**) ChIP-qPCR signal for facultative heterochromatin mark H3K27me3. (**C**) qPCR signal revealing chromatin accessibility to nucleases.

Heterochromatin is more compact than actively transcribed chromatin. We determined chromatin accessibility by treating isolated chromatin with a panel of nucleases and amplifying the resulting fragments with qPCR. As expected from the heterochromatin results, in time, the proviral chromatin was compacted; its accessibility declined over time (Fig 5C).

### Loss of proviral H3K27ac, but heterochromatin maintained during activation

Next, we asked how the proviral chromatin landscape changed during T-cell activation. Cells at 30 dpi were TCR-stimulated (αCD3-CD28) or mock-treated. After 48 h, the chromatin and DNA were isolated, and we interrogated H3K27ac, H3K27me3, and H3K9me3 levels with ChIP-qPCR (Fig 6A). The purified genomic DNA revealed that 80% of proviruses remained detectable after activation; i.e., through proliferation and cell death, only 20% of provirus was lost after one round of activation. As expected, the stable heterochromatin marks, H3K9me3 and H3K27me3, remained unchanged in the core of the provirus. However, the levels of the active enhancer mark, H3K27ac, dropped significantly at the four positions throughout the provirus (p<0.05). This would be expected, if the H3K27ac was located on chromatin that was targeted for reactivation. Interestingly, the levels of H3K27ac dropped below the levels of the input DNA. This finding suggested that two mechanisms must be at work; one was the loss of H3K27ac labeled proviruses and the second was the active removal of the H3K27ac-mark from the proviruses. Removal of H3K27ac is associated with chromatin transitioning from an active to a poised state [53].

**Fig 6.**
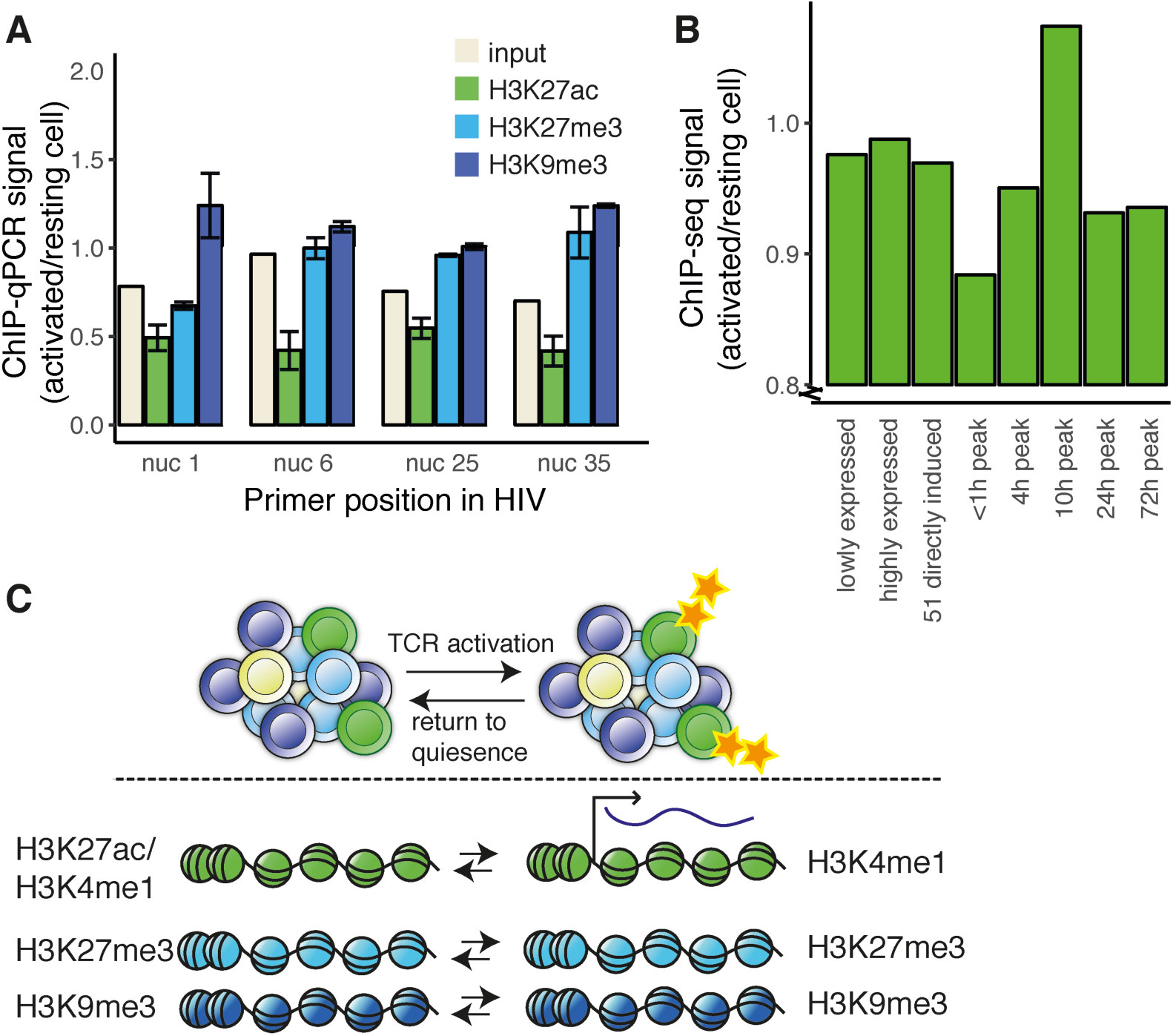
Chromatin modulation following TCR activation. (**A**) H3K27ac, H3K27me3 and H3K9me3 ChIP over the HIV-1 provirus. DNA and chromatin isolated 48h after T-cell activation (or mock-treatment) of Bcl2-donor cells. Experiments performed in triplicate from a single donor, error bars represent s.e.m..(**B**) H3K27ac ChIP seq of the same sample as in A. Ratios for ChIP signal activated/resting cells averaged over eight previously published gene sets. (**C**) Graphical summary of the model with a heterogeneous HIV-1 reservoir of functionally intact latent proviruses and selective and transient activation of the provirus following T-cell activation.

We subjected the same H3K27ac ChIP sample to genome-wide sequencing. The sequencing reads from the provirus were too few to support firm conclusions, but we could determine H3K27ac patterns associated with cellular genes. First we confirmed technical soundness by comparison to ENCODE datasets of H3K27ac ChIP in primary CD4 T-cells. We constructed probes of 2kb centered on the 5’ site of all genes. The large 2kb window was chosen as the H3K27ac mark spanned the HIV-1 provirus. The Pearson’s correlations between signals in the datasets were 0.52, with a clear positive trend (Fig S5). During TCR activation, the H3K27ac showed no genome-wide changes. We interrogated lists of genes affected at the transcriptional level when primary T cells were exposed to TCR activation [54] (Table S3). We examined H3K27ac levels at the 2kb-defined probes prior and post activation. Genes with consistently low expression, consistently high expression, or the 51 genes most significantly induced (at 48 h) showed no change in H3K27ac levels with activation (Fig 6B). However, five gene sets showed changes in gene expression, with transcriptional peaks between 0 h and 72 h; these displayed variable degrees of changes in H3K27ac levels. One gene set showed the highest expression within the first hour after activation, but then became repressed. This gene set showed a down-regulation of H3K27ac at 48 h after TCR activation. As we just demonstrated, this was the expression profile observed for latent provirus. Consequently, this result indicated that the latent, reactivatable HIV-1 provirus belonged to the set of genes that were transiently affected by T-cell activation, but then repressed.

## Discussion

This study demonstrated that the reservoir of latently HIV-1 infected cells comprised proviruses with heterogeneous modes of silencing, with highly different reactivation potential. From a functional cure perspective, the part of the reservoir that requires elimination is the fraction that can be reactivated, because that is the only compartment capable of a *de novo* spread of virus. Our data suggested that these reactivatable proviruses were marked with the enhancer mark H3K27ac, in quiescent T-cells. This finding was supported by our previous study, which showed that reactivatable proviruses were largely integrated into enhancer regions marked with H3K4me1 [20]. H3K4me1 and H3K27ac have mainly overlapping distributions, but H3K4me1 is relatively stable, and H3K27ac fluctuates with expression [53]. Here, we propose a model (Fig 6C) where, upon T-cell activation, the proviruses marked by either H3K9me3 or H3K27me3 heterochromatin marks remain unaffected, consistent with heterochromatin tending to be stable during perturbations [36,55-57] In contrast, upon T-cell activation, proviruses with enhancer marks, H3K27ac and H3K4me1, are expressed, which results in the production of viral particles. HIV-1 proviruses located at the nuclear pore are associated with enhancer chromatin [24], which allows rapid nuclear export. After activation, the provirus loses its H3K27ac mark, and it is actively re-silenced, consistent with the genome-wide trend of early repressed genes (Fig 6B). The loss of H3K27ac during transcription activation may be a consequence of histone replacement [57].

Although we have long understood the positive feedback loop mechanism driven by Tat [38], we lack knowledge of mechanisms that actively silence the provirus. However, recent studies have revealed negative HIV-1 feedback loops that might rely on RNA precursor export [58] or histone modifications through the arginine methyltransferase CARM1 [59]. Interestingly, H3K27ac promotes CARM1-mediated HIV-1 latency. This association suggested the hypothesis that a rapid transient pulse of HIV-1 followed by programmed silencing may reseed the infection, but prevent cytopathic effects of the virus and prevent immunological detection.

The evidence presented here has further challenged the notion that promoter-proximal RNAPII pausing is the major rate-limiting step in HIV-1 expression. Instead, we have confirmed that, in latently infected cells, the appearance of mature HIV-1 transcripts was blocked after the RNAPII initiation step. The observation that promoter-proximal RNAPII was elongating in latently infected cells (Fig 4B) suggested that the P-TEFb-subunit, CTD ser2-kinase, CDK9, was also present during latency. P-TEFb interacts with the BET protein BRD4, and BRD4 is found at the HIV-1 promoter, where it performs a silencing function [39]. BRD4 recognizes H3K27ac, which explains the presence of this mark at the inactive, but poised, provirus. Silencing BRD4 and activating Tat compete for P-TEFb binding [60,61]; this competition implied that sufficient levels of Tat, produced from multiply spliced transcripts, would be required to overcome this post-elongation block.

Current cure efforts (using the “shock-and-kill” approach) with LRAs have managed to increase viral RNA levels in patients, but this approach has shown no or very limited effect on the reservoir size. A previous model has shown that a medication-free sustained remission (“cure”) in 50% of HIV-1 positive individuals would require reducing the reservoir by >4 log units [62]. However, the heterogeneous reservoir mainly consists of non-functional proviruses; therefore, we need to estimate the fraction of the reservoir with reactivation potential. The provirus can become non-functional as a consequence of mutations, but also from other, insurmountable challenges, which result from epigenetic or transcriptional obstacles. Thus, either PCR-amplifying short provirus regions or sequencing the intact provirus provides an overestimation of the functional reservoir [63]. Furthermore, some transcription occurs over the provirus without producing mature functional transcripts (Fig 1A) [40]; thus, HIV-1 detection with RNA *in situ* hybridization would also overestimate the functional reservoir. Here, we have stressed the establishment and maintenance of latency through epigenetic and direct transcriptional processes; however, other factors also affect latency. These include transcriptional interference; limiting concentrations of transcription factors–notably NFkB, NFAT, and AP1; and the host metabolism [64]. In addition, post-transcription failure to produce viruses can be a result of RNA processing or variations in the functions of viral proteins [40,63].

In summary, our findings pinpointed some discrepancies among model systems that have hampered our understanding of HIV-1 latency. In addition, we have presented a way to identify the activatable fraction of the heterogeneous latent HIV-1 reservoir. By manipulating the activity of the latent reservoir, the disease burden may be reduced in individuals living with HIV-1.

## Materials and Methods

### Ethics Statement

This study was approved by the Ethics Committee (Regionala Etikprövningsnämnden Stockholm, Reg#2017/1138-31 and Reg#2018/102-31), and written informed consent was obtained from all subjects. The data were analyzed anonymously.

### Human samples

Buffy coats from 450-ml blood samples drawn from healthy donors were provided by the Karolinska Universitetslaboratoriet. The samples were anonymized before arrival. Patient samples were obtained from the HIV unit at Department of Infectious Diseases, Karolinska University Hospital.

### Cell culture

Bcl2 model cells were generated as previously described [46]. Peripheral blood mononuclear cells (PBMCs) were purified on Ficoll-paque PLUS (GE Healthcare, Cat#17-1440-02). CD4^+^ T lymphocytes were extracted (Milentyi Biotec Cat#130-096-533) by negative selection. Resting cells were kept in RPMI 1640 medium (Hyclone, Cat# SH30096_01), 10% FBS (Life Technologies, Cat# 10270-106), 1% Glutamax (Life Technologies, Cat# 35050), 1% Penicillin-streptomycin (Life Technologies, Cat# 15140-122). For active growth conditions, media was supplemented with human interleukin-2 IS (Miltenyi Biotec, Cat#130-097-742; Lot#5170516373) final concentration 100U/ml and 5% T-cell conditioned media, according to the protocol.

### Virus production

EB-FLV (containing *bcl2*), pNL4-3-Δ6-drEGFP (reporter HIV-1), pHelper [47], and pMD2.G (VSV-g) (Addgene, Cat#12259) plasmids were purified with Plasmid Plus Maxi Kit (Qiagen, Cat# 12963). 293T cells (ATCC, CRL-3216; CVCL_0063) grown in DMEM media (Hyclone, Cat# SH30022_01) were transfected with Lipofectamine LTX with PLUS reagent (ThermoFisher, Cat# 15338100), and, after an additional 48 h, supernatants were harvested. We tested the functional infectivity of NL4-3-Δ6-drEGFP by transducing 293T cells (American Type Culture Collection, Cat# CRL-3216) with the viral particles. After 48 h, we measured GFP signals with flow cytometry. We determined virus titers by the HIV-1 p24 ELISA Assay (XpressBio, Cat# Cat#XB-1000). Virus-containing supernatant was concentrated with LentiXconcentrator (Clontech, Cat# 631231).

### Infection

Prior to infection, cells were activated for 72 h in media with 1 μg/ml anti-CD28 (BD, Cat# 555725) in 6-well plates coated for 1 h at 37°C with 10 μg/ml anti-CD3 (BD, Cat# 555336). Cells where then spinoculated (2h at 1,200*g* 25°C) with pseudotyped EB-FLV or NL4-3-Δ6-drEGFP at a concentration of 250 ng p24 per 1×10^6^ cells.

### Chemicals to induce proviral activation

Cells were exposed to latency-reversal agents for 48 h (or as indicated). Drugs and chemicals used were phorbol 12-myristate 13-acetate (Sigma-Aldrich, Cat# 79346) final concentration 50 ng/ml, ionomycin (Sigma-Aldrich, Cat# I0634; Lot#106M4015V) final concentration 1 nM, panobinostat (Cayman Chemicals, Cat# CAYM13280) final concentration 30 nM, JQ1 (Cayman Chemicals, Cat#CAYM11187) final concentration 100nM, bryostatin (Biovision, Cat# BIOV2513) final concentration 10nM. For all treatments, raltegravir (Sigma-Aldrich, Cat# CDS023737) was added to the medium at final concentration 2 μM.

### Flow cytometry

Cells were stained with mouse anti-human CD4 PE-Cy5 (clone RPA-T4, BD 561004); CD25 APC (clone M-A251, BD 560987); CD69 PE-Cy7 (clone FN50, 557745), LIVE/DEAD Fixable Violet Dead Cell Stain (ThermoFisher, Cat# L34955), and fixed in 2% folmaldehyde for 30 min. Flow analysis was performed on a CytoFLEX S (Beckman Coulter). Individual flow droplets were gated for lymphocytes, viability, and singlets. Data was analyzed by Flowjo 10.1 (Tree Star).

### Droplet digital PCR (ddPCR)

ddPCR was performed with the QX200 Droplet Digital qPCR System (Bio-Rad). Samples were tested in duplicate, and each reaction consisted of a 20-µl solution containing 10 µl Supermix for Probes without dUTP (Bio-Rad, Berkeley, CA, USA), 900 nM primers, 250 nM probe (labeled with HEX or FAM), and 5 µl undiluted RT product or 500 ng cellular DNA (fragmented with a QIAshredder column). Emulsified PCR reactions were performed with a C1000 Touch thermal cycler (Bio-Rad), with the following protocol: 95°C for 10 min, followed by 40 cycles of 94°C for 30 s and 60°C for 60 s, and a final droplet cure step of 10 min at 98°C. Each well was then read with the QX200 Droplet Reader (Bio-Rad). Droplets were analyzed with QuantaSoft, version 1.5 (Bio-Rad), software in the absolute quantification mode. When replicates were used, the percentage of mutant fractional abundance was extracted as merged samples. For visualization, we used the “twoddpcr” Bioconductor/R package [65]. Nucleotide numbers are set according to the coordinates of the reference Human immunodeficiency virus type 1 (HXB2; K03555)

### Measurement of intracellular HIV-1 transcripts

Total cellular RNA was isolated from 1×10^6^ latently infected Bcl2-transduced cells with RNeasy Mini Plus Kit (Qiagen, Cat# 74134). RNA (500ng) was used directly in reactions with SuperScript III Reverse Transcriptase (Invitrogen, Cat#11752-050), primed by random hexamers (ThermoFisher, Cat#S0142). Reactions were incubated at 25°C for 10 min, followed by 50°C for 30 min. Reactions were terminated at 85°C for 5 min followed by incubation on ice. Subsequently, 2U/reaction of *E.coli* RNAse H (Invitrogen, Cat#18021-014) was added and tubes were left at 37°C for 20 min, after which they were stored at −20°C. cDNA was specifically quantitated at specific positions with ddPCR (Table S3).

### Chromatin immunoprecipitation-PCR

Prior to chromatin extraction, viable cells were isolated using Ficoll density separation (300*g* for 10 min at room temperature). ChIP-qPCR was performed using the iDeal ChIP-qPCR Kit (Diagenode, Cat# C01010180). Each ChIP reaction was performed on 1 × 10^6^ cells. Sonication was performed at 30s in eight cycles (Bioruptor Pico, Diagenode, Cat# B01060010). ChIP antibodies were targeting H3 (Abcam, Cat# ab1791), H3K4me3 (Diagenode, Cat# C15410030), H3K9me3 (Abcam, Cat# ab8898), H3K27me3 (Diagenode, Cat#C15410069), H3K27ac (Abcam, Cat# ab4729), RNAPII-ser2ph (Diagenonde, Cat# C15200005), RNAPII-ser5ph (Diagenode, Cat# C15200007), IgG (Diagenode, Cat# C15410206). Primer sequences are shown in Table S3. PCR reactions were performed with Powerup Sybr green master mix (2x) (ThermoFisher, Cat#A25742) using 40 cycles on an Applied Biosystems 7500 Fast Real-Time PCR System (ThermoFisher).

### Chromatin accessibility

Nuclease accessibility was evaluated through the Chromatin Accessibility Assay Kit (Abcam, Cat# ab185901) according to manufacturer’s instructions. Per reaction, 0.5 × 10^6^ cells were used.

### Sequencing

DNA samples were quantified with Qubit dsDNA HS Assay kit (ThermoFisher, Cat# Q32851) and libraries were prepared using NEBNext Ultra II DNA library kit they were sequenced on an Illumina Hiseq 2000 (50 cycles, single-end sequencing, 50 bases) at the BEA facility (Huddinge, Sweden), according to the manufacturer’s instructions. Raw data from the Hiseq (fastq files) were aligned to the hg19 genome assembly with the Bowtie2 program (version 2.0.6), set to the default parameters. Resulting sam files were converted to bam files using Samtools version 1.4. Bam files were imported into SeqMonk version 0.33.0 where 2kb probes were constructed around the 5’ position of all 40,147 genes of the GRCh37 assembly. Probes were quantitated with ‘Read Count Quantitation’ using ‘All Reads’ correcting for total count per million reads, duplicates were ignored.

RNA-seq (mRNA) data from primary CD4 cells were collected from GSM669617 (GEO). H3K27ac ChIP-seq data for comparison were collected from ENCFF618IUD and ENCFF862SKP (Encode).

### Data availability

The H3K27ac ChIP-seq data have been deposited in the GEO database under ID GSE121055.

## Acknowledgments

The plasmids, pCM6 and pC-Help, were gifts from Robert Silicano, and the pMD2.G plasmid was a gift from Didier Trono (Addgene plasmid # 12259). We thank Andreas Lennartsson for critical reading of the manuscript. We would like to acknowledge the core facilities MedH Core Flow Cytometry facility (Karolinska Institutet) for providing cell analysis services, and BEA, Bioinformatics and Expression Analysis (Karolinska Institutet) for providing sequencing services.

## Supplementary material

**Fig S1:**
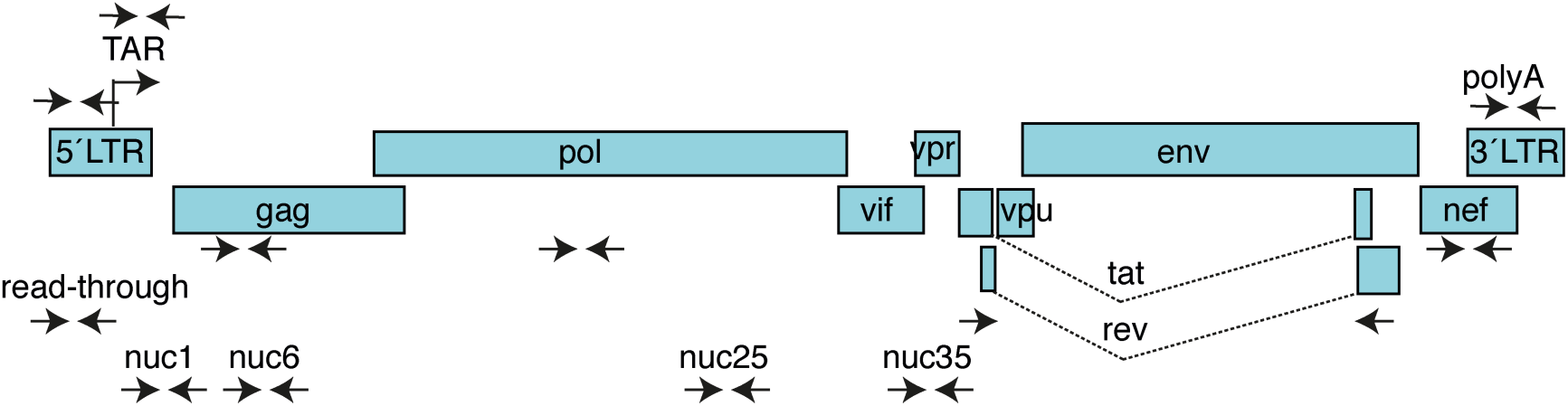
Primer positions relative to the HIV-1 provirus. Viral proteins depicted with blue bars.

**Fig S2:**
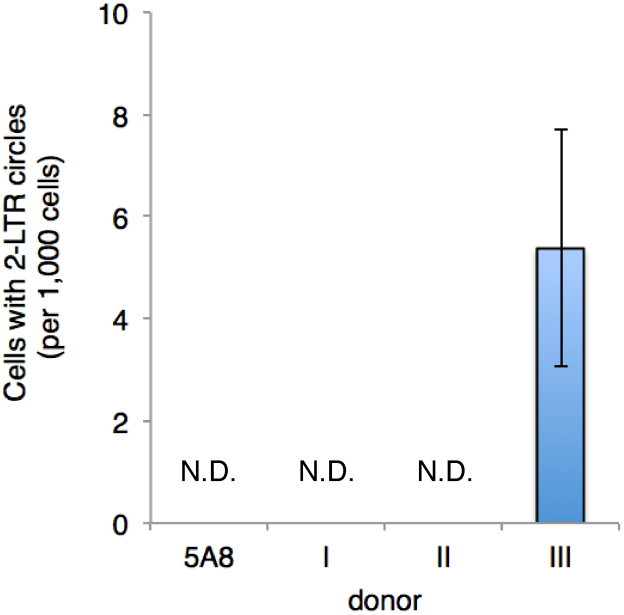
Quantification of 2-LTR circles. RNA was isolated at 3 dpi. Resulting cDNA was tested using Taqman PCR with probes recognizing 2-LTR circles and host *rpp30* for normalization. N.D. not detectable.

**Fig S3:**
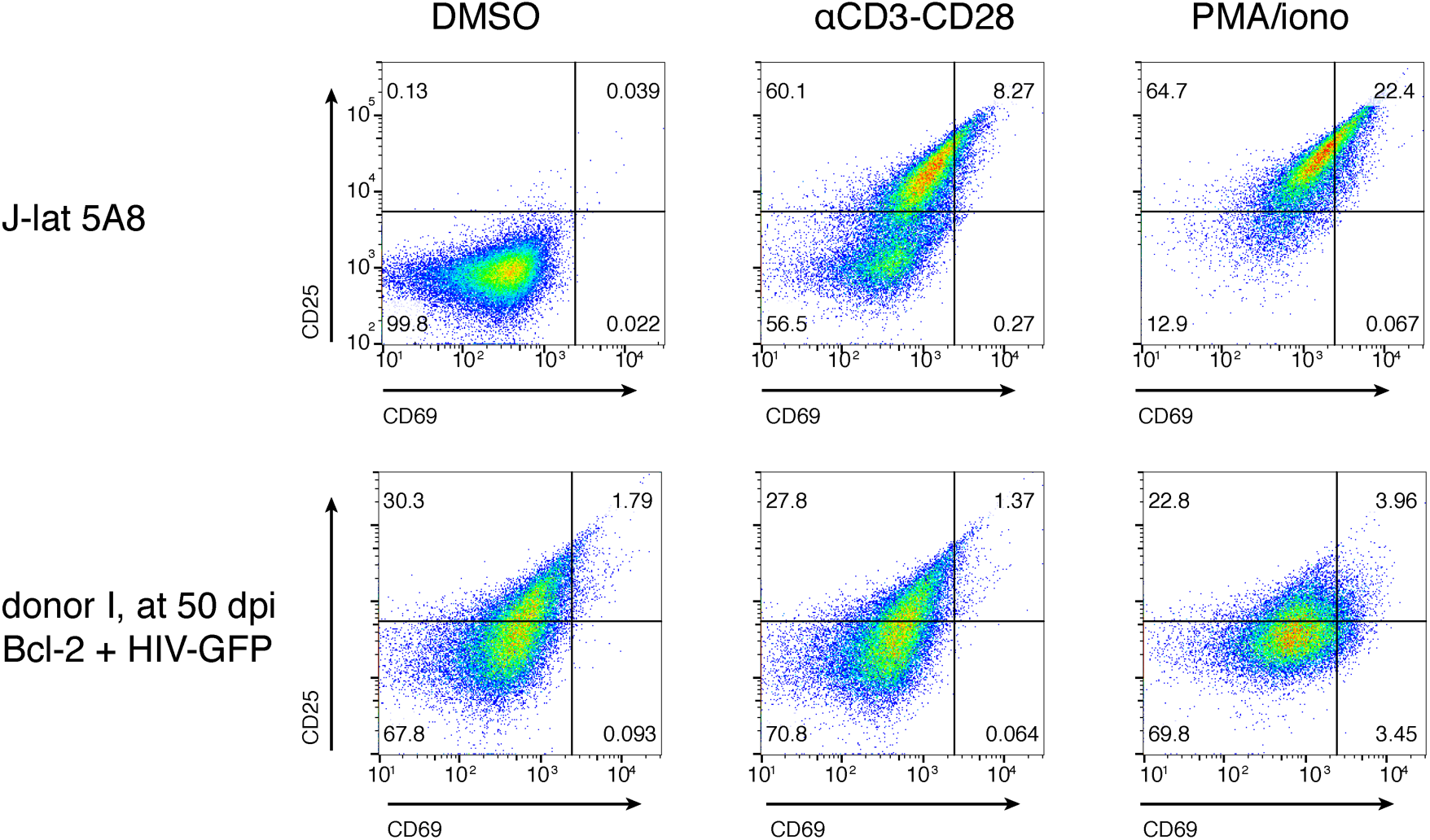
T-cell activation. Bcl2 cells with HIV-1-GFP at 50 dpi and J-lat 5A8 cells were exposed to DMSO, antibodies against CD3 and CD28, or PMA/ionomycin for 48 h prior to flow cytometry analysis using labeled antibodies against surface markers CD25 and CD69.

**Fig S4:**
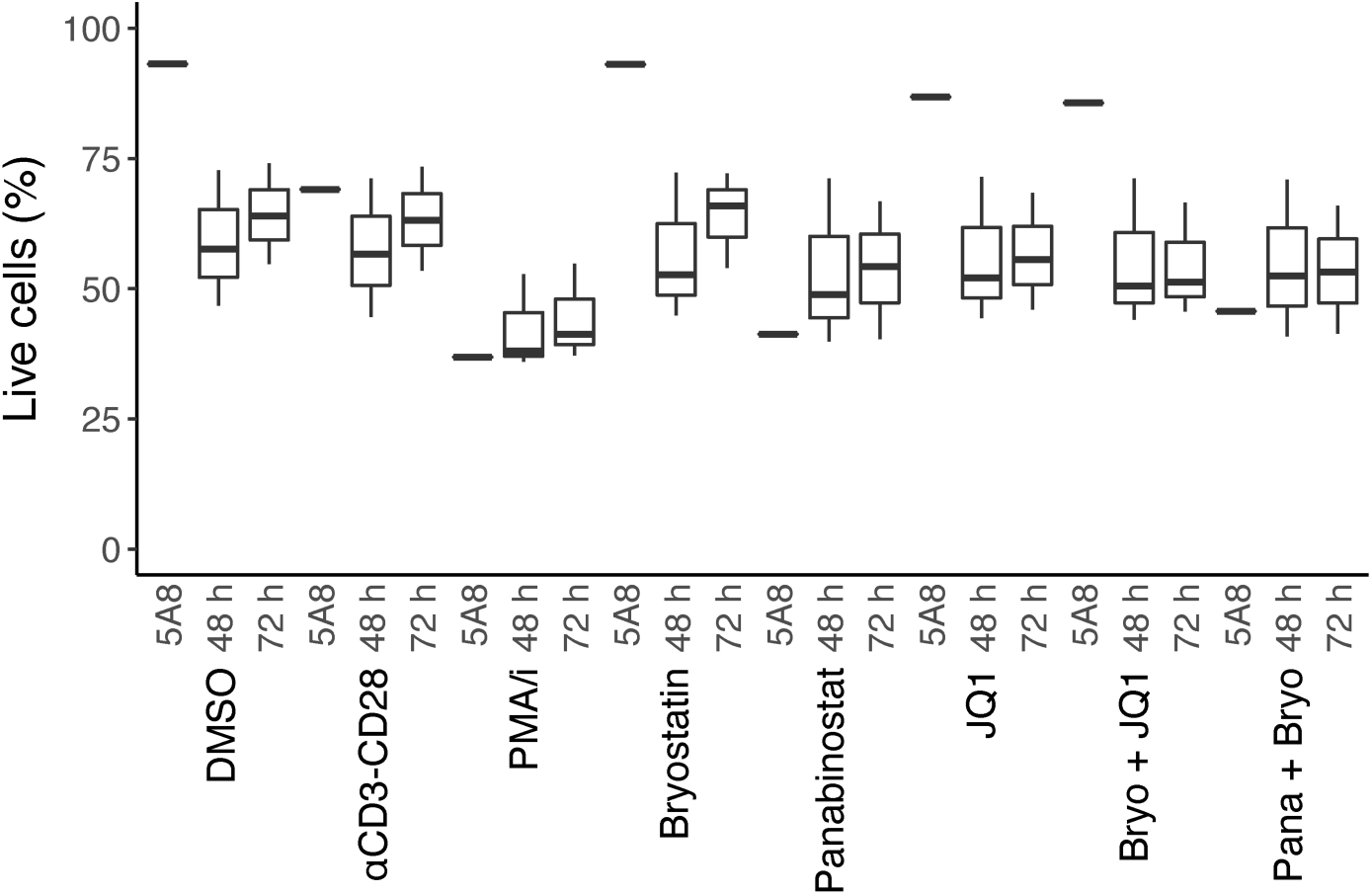
Cell viability after drug exposure. Boxplot showing the cell viability as determined by membrane integrity through LIVE/DEAD staining and flow cytometry. HIV-1 infected Bcl2 model cells from healthy donors (*n*=3) were exposed to drugs for 48h and 72h. J-lat clone 5A8 was used as control.

**Fig S5:**
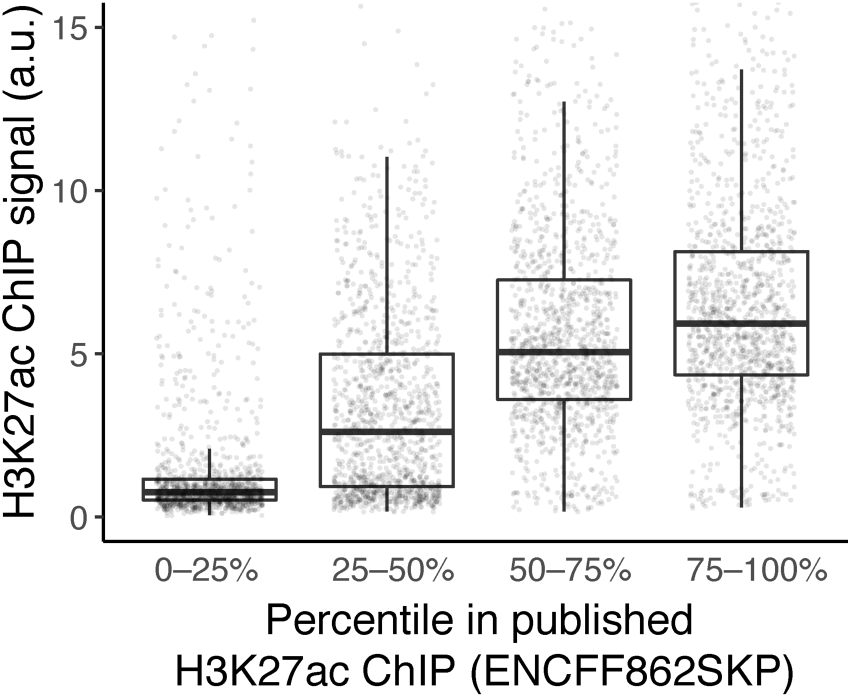
Comparison between H3K27ac ChIP in HIV-1-GFP infected Bcl2-model cells from healthy donor I and ENCODE dataset. Boxplot showing the H3K27ac ChIP signals (resting CD4 T-cells) calculated in 2kb-probes centered around the start of genes. Published ChIP data (ENCODE ENCFF862SKP) were processed in the same way and grouped in quartiles. All individual data points are shown.

**Table S1:** Patient characteristics.

| Patient ID | Sex | HIV subtype | Year HIV+ | Earliest CD4 count |  | Earliest Plasma HIV-1 RNA |  | Times before sampling |  |  | Values at sampling |  |  |
| --- | --- | --- | --- | --- | --- | --- | --- | --- | --- | --- | --- | --- | --- |
| | | | | Year | Value (cells/μL) | Year | Value (copies/mL) | Time on ART | Time with VL<50 (copies/mL) | Lowest CD4 detected (cells/μL) | CD4 count (cells/μL) | Viral load (copies/mL) | ART regime |
| 1 | F | HIV-1C | 2013 | 2013 | 560 | 2013 | 295 | 1.8 yrs<sup>1</sup> | 1.8 yrs | 520 | 620 | <20 | 3TC/ABC/DTG |
| 2 | F | HIV-1C | 1999 | 1999 | 420 | 1999 | 20,600 | 9 yrs | 8 yrs<sup>2</sup> | 150 | 640 | <20 | TAF/FTC/EVG/cobicistat |
| 3 | F | CRF | 2014 | 2014 | 210 | 2014 | 26,500 | 4 yrs | 3.8 yrs | 200 | 620 | <20 | 3TC/ABC/EFV |
| 4 | F | HIV-1A | 1989 | 1989 | 210 | 1996 | 499 | 18 yrs<sup>1</sup> | 16 yrs | 140 | 390 | <20 | RPV/DRV/r |
| 5 | M | ND | 1997 | 1997 | 173 | 1997 | 136,000 | 20 yrs | 20 yrs | 231 | 800 | 36 | 3TC/ABC/EFV |

Abbreviations: 3TC, lamivudine; ABC, abacavir; CRF, circle recombinant form; DRV, darunavir; DTG, dolutegravir; EFV, efavirenz; EVG, elvitegravir; F, female; FTC, emtricitabine; M, male; ND, Not determined; RPV, rilpivirine; r, ritonavir-boosted; TAF, tenofovir alafenamide.
<sup>1</sup> Patient 1 started treatment 5 months 2014 but it was interrupted and started again in Sep 2016. Patient 4 started treatment 1996-1999 but was interrupted between 1999-2000
<sup>2</sup> One blip in Viral Load in 2014

**Table S2: H3K27ac ChIP signal over 2kb probes of genes.** Quantification of sequencing reads overlapping 40,147 probes designed around the 5’region of genes in the GRCh37 assembly. Columns indicating belonging to a gene set from Fig. 6B.

*Large table in accompanying excel file.*

**Table S3:**
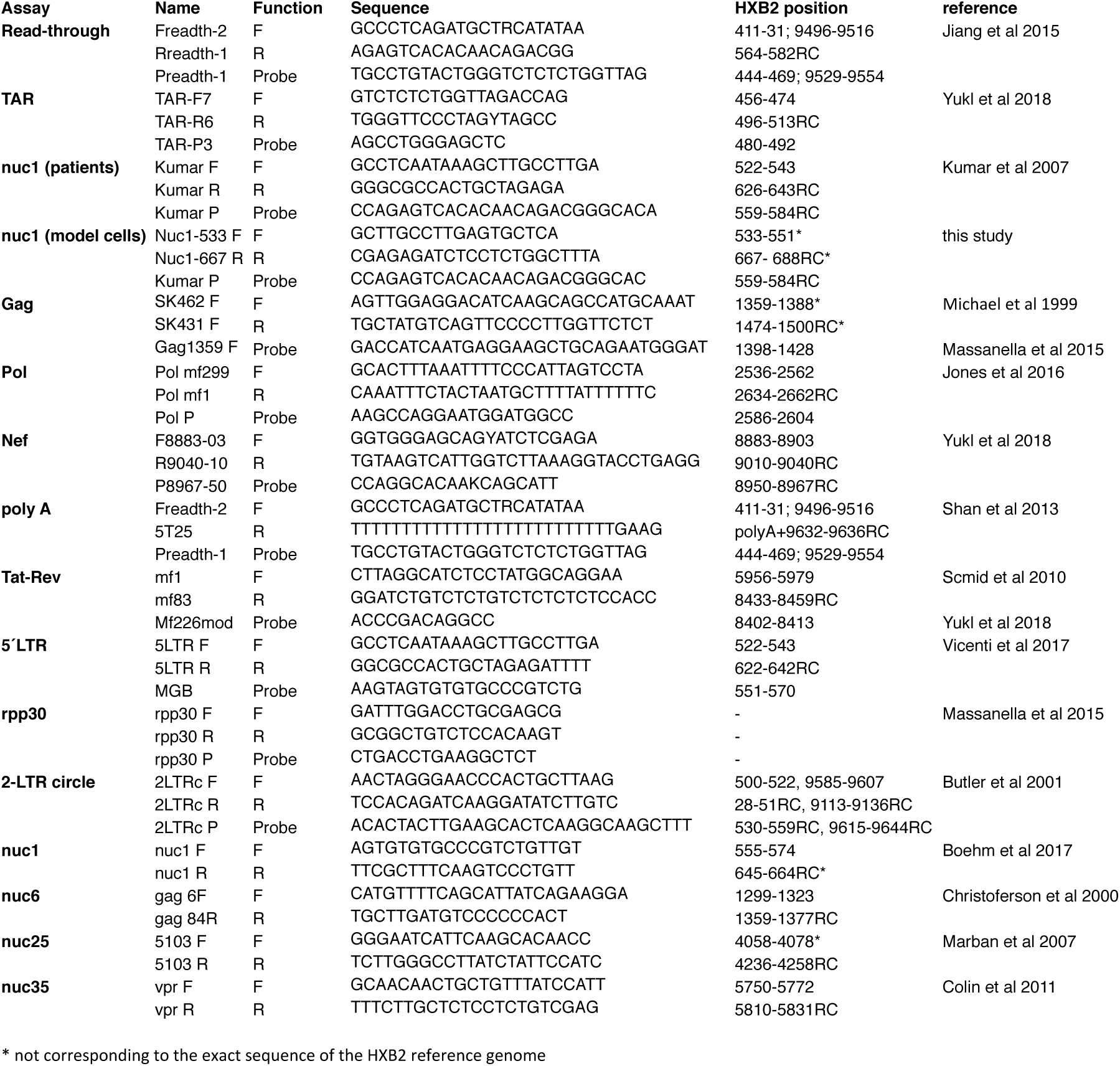
Primer and probe sequences. Positions are given relative to the HXB2 reference genome.

